# Phytochemical screening, analgesic, anti-pyretic and antibacterial potentials of *Litsea glutinosa* (L) leaves extracts *in vivo* and *in vitro* technique

**DOI:** 10.1101/2024.08.21.609048

**Authors:** Zubair Khalid Labu, Samira Karim, Md. Tarekur Rahman, Md. Imran Hossain, Md Shakil

**Affiliations:** Department of Pharmacy, World University of Bangladesh, Dhaka, Bangladesh

**Keywords:** *Litsea glutinosa*, Phytochemical screening, analgesic, antipyretic, antibacterial activities

## Abstract

**Background:** *Litsea glutinosa* leaves (LG) have been used as traditional medicine to treat diseases since ancient times. This study aimed to investigate the cold methanol extract of LG for its analgesic, antipyretic and antibacterial activities, along with preliminary phytochemical screening and acute toxicity test.

**Methods:** In this study, we first investigated the major phytochemicals group present applying modern chromatographic technique HPLC in addition to conventional methods for Phyto screening in cold methanol extracts LG leaves. Both methods simultaneously demonstrated major bioactive compounds found in the extract were phenol, flavonoid. The purposive efficacy and toxicity were then assessed through preclinical testing

**Results:** In hot plate method, the highest pain inhibitory activity was found at a dose of 500 mg/kg of crude extract (3.37± 0.31sec) which differed significantly (P <0.01 and P <0.001) with that of the standard drug morphine (6.47± 0.23 sec). The extract significantly prolonged reaction latency to thermal-induced pain in hotplate model. Analgesic activity at 500 mg/kg, LG extract produced a 70% suppression of writhing in mice, which was statistically significant (*p<0.001*) compared to standard morphine’s (77.5%) inhibition. In antipyretic activity assay, the crude extract showed notable reduction in body temperature (36.17 ± 0.32°C) at dose of 300 mg/kg-body weight, when the standard (at dose 100 mg/kg-body weight) exerted (36.32 ± 0.67°C) after 3 h of administration. In antibacterial studies, results showed that inhibition of bacterial growth at 400 μg dose of each extract clearly inhibited growth of bacteria from 11 to 22 mm. The extractives carbon tetrachloride fraction, chloroform soluble fraction, ethyl acetate fraction demonstrated notably greater inhibitory zone widths (p < 0.05) against tested strains.

**Conclusion:** Overall, the cold methanol extract of LG leaves demonstrates the therapeutic potential in preclinical settings.

## 1. Introduction

In recent times, plant-based natural compounds have gained global prominence as complementary and alternative medicine, significantly contributing to the enhancement of health and well-being. Many widely used pharmaceuticals, such as aspirin, digoxin, morphine, ephedrine, quinine, tubocurarine, and reserpine, have their origins in medicinal plants. Phenols and flavonoids are particularly valued phytoconstituents due to their hydroxyl groups, which enable them to decompose peroxides, repair oxidative damage, and quench singlet and triplet oxygen [1]. The traditional knowledge of medicinal plants and their uses by indigenous cultures is not only vital for preserving cultural traditions and biodiversity but also essential for community healthcare and modern drug development. Synthetic tropical therapies often come with numerous side effects and are costly, making them inaccessible to many. To address this issue, plants available in the vicinity are often used without scientific validation. The use of higher plants and their extracts to treat infections is a long-established practice, with herbal medicines gaining popularity for their cost-effectiveness and eco-friendliness [2].

*L. glutinosa*, belonging to the Lauraceae family, is a well-known evergreen species found in the forests of Chittagong and Sylhet districts in Bangladesh. It is occasionally cultivated in various parts of the country. The leaves, known for their mucilaginous properties, are used as antispasmodic, emollient, and poultice treatments. They are also employed in treating diarrhea, dysentery, wounds, and bruises [3]. The leaves have been reported for their use in treating the spontaneous and excessive flow of semen in young boys [4]. Additionally, the leaves extract exhibits antibacterial, analgesic and cardiovascular activities [5]. The berries produce oil that some tribal practitioners use to treat rheumatism. Common constituents of this species include tannin, β-sitosterol, and actinodaphnine, along with other compounds such as Boldine, norboldine, laurotetanine, n-methyllaurotetanine, n-methyl actinodaphnine, quercetin, sebiferine, and litseferine [6].Pain, an unpleasant sensory and emotional experience associated with actual or potential tissue damage, is relieved by analgesic compounds that act on the central nervous system or peripheral pain mechanisms without significantly altering consciousness. Analgesics are typically used when the noxious stimulus cannot be removed or as an adjunct to more etiological pain treatments [7,8]. Inflammation, the response of living tissues to injury, involves a complex array of enzyme activation, mediator release, fluid extravasation, cell migration, tissue breakdown, and repair. Non-steroidal anti-inflammatory drugs (NSAIDs) are frequently prescribed due to their effectiveness in treating pain, fever, inflammation, and rheumatic disorders. However, their use is linked to adverse effects on the digestive tract, ranging from dyspeptic symptoms, gastrointestinal erosions, and peptic ulcers to more severe complications such as bleeding or perforation [9]. Thus, developing new anti-inflammatory drugs with fewer side effects remains crucial, and natural products like medicinal plants could lead to discovering new, safer anti-inflammatory agents [10].

Pyrexia, or fever, often results from infections, tissue damage, inflammation, graft rejection, malignancy, or other diseased states. The body’s natural defense mechanism raises the temperature to create an environment where infectious agents or damaged tissue cannot survive. Typically, infected or damaged tissue initiates the increased formation of pro-inflammatory mediators (cytokines like interleukin 1β, α, β, and TNF-α), which enhance the synthesis of prostaglandin E2 (PGE2) near the preoptic hypothalamus area, triggering the hypothalamus to elevate body temperature [11]. Most antipyretic drugs prevent or inhibit COX-2 expression to reduce elevated body temperature by inhibiting PGE2 biosynthesis. However, synthetic agents irreversibly inhibit COX-2 with high selectivity, which can be toxic to hepatic cells, glomeruli, the brain cortex, and heart muscles, whereas natural COX-2 inhibitors usually have lower selectivity and fewer side effects [12]. *L. glutinosa* was chosen for this study due to its availability in Bangladesh and its traditional use in rural areas for various treatments. To date, no investigations have been conducted on this plant native to Bangladesh. Our primary aim was to evaluate the in antipyretic, analgesic activities and antibacterial properties of the plant leaves to validate their traditional uses.

## 2. Materials and Methods

### 2.1 Plant sample collection and accurate documentation

Plant material was collected from Mirjapur village in the Tangail district of Bangladesh (geographical coordinates: 24.05° N, 89.92 ° E), July 2023. where the plant leaves were collected. After completing the identification of plant leaves by Bangladesh National Herbarium’s taxonomist followed by provided taxonomical accession number # 354079 and with all necessary permissions obtained to continue our study.

### 2.2 Chemicals and instruments

Analytically graded chemicals were used throughout the study, including methanol (liquid chromatography grade, ≥99.8%), ethyl acetate (≥99.9% GC), pet-ether (≥80%), chloroform (≥99% ACS Reagent Grade), carbon tetrachloride (≥99.9%), gallic acid (98%), catechin (≥99.8%), Folin-Ciocalteu reagent (standard reagent grade), and aluminum chloride (anhydrous sublimed, ≥99.8%), all provided by Science Park Chemicals Ltd., Bangladesh. The instruments used were calibrated according to standard protocols to minimize experimental errors. The list of instruments includes:

UV-Vis spectrophotometer (Model: UV-1700 series), Casio Digital Stopwatch (Model: HS-70W-1DF), Mechanical dryer (CG-CG23KW-150KW), Digital Analytical Balance (Model: PS. P3.310), Thermostat water bath (Model: HHW21.420AII), Autoclave (Model: DSX-280KB), Rotary evaporator (Model: PGB002), Vortex mixer (Model: VM-11), Sonicator (Model: ULP-3000), HPLC (LC-2060 3D HPLC system, a Shimadzu model)

### 2.3 Drying and grinding of plant materials

After being separated from unwanted leaves or materials, the gathered plant components (leaves) were cleaned with water to remove any remaining dirt. After being sun-dried for a week, they were dried in a mechanical dryer. Before analysis could begin, the plant parts were placed in an airtight container, mechanically processed into a coarse powder, and maintained in a cool, dry, and dark place to ensure optimal storage and preservation.

### 2.4 Plant Material Extraction

400 gm of the substance as powder form was put in a spotless, tinted glass container with a flat bottom and were submerged in 1500 ml of methanol at 25°C temperature. The container and its contents were securely closed to prevent air from entering, and they were kept for ten days with sporadic shaking and stirring to improve extraction. After decanting the extract through cotton, Whatman no. 1 filter paper was used to filter it. Using a rotary evaporator, the final filtrates were concentrated at 40°C [13]. It created a sticky, greenish-black concentration. It was determined that the sticky concentration was crude methanol extract. Eleven grams of the crude methanolic extract were dissolved in 90% methanol using the Kupchan technique, and the mixture was then divided among organic solvents with different polarity [14]. Complete evaporation of the fractions, the following fractions were produced: ethyl acetate fraction (ESF, 4.2gm), pet-ether fraction (PSF, 1gm), chloroform fraction (CSF, 5gm), carbon tetrachloride fraction (CTF, 1.5gm), methyl soluble fraction (MSF, 2gm) and the remaining aqueous fraction (AQF, 0.8 gm). The resulting fractions were kept in cold place for biological analysis.

### 2.5 Phytochemical screenings

A confirmatory qualitative phytochemical screening was performed on all crude extracts with a slight modification of Sofowara’s standard protocol for phytochemicals in the crude extracts [15, 16]. The main classes of compounds obtained were various organic soluble fractions, for example, glycosides, resins, phenols, alkaloids, saponins, steroids, and flavonoids.

### 2.6 High performance liquid chromatography (HPLC)

HPLC stands as the well-known method for the chemical standardization of plant extracts. In this research study, gallic acid served as the standard for phenolic compound identification, while catechin functioned as the standard for identifying flavonoid compounds. The HPLC analysis employed a LC-2060 3D HPLC system, a Shimadzu model from Japan. The C18 column (ID: 5-micron x 100 Å) was used as the stationary phase, which gives a larger surface area the mobile phase has to travel across. The HPLC system incorporated a data detector A - Ch1. The mobile phase, operating in an isocratic gradient system, comprised 30% acetonitrile and 70% water. Before use, the mobile phase underwent filtration through a 0.45 μm filter paper and degas by sonication. The flow rate was selected at 1 mL/min. In this case, methanol was adjusted to 0.75 mL/min and distilled water was adjusted to 0.25 mL/min and the HPLC column was maintained at a temperature of 45°C. Sample injection involved a volume of 20 μL, run time for the sample of 10 min and standard 5 min.

### 2.7 Estimation of total phenolic content (TPC)

With a few minor modifications, the recommended methodology was applied in the experiment to determine TPC using the Folin-Ciocalteu reagent [ 17]. Two milliliters of crude methanolic extract and all of its organic soluble components were liquefied with two milliliters of distilled water to achieve the final concentration of one milligram per milliliter. A mixture of 0.5 mL of extractives (1 mg/mL) and 2mL of diluted Folin-Ciocalteu (previously diluted 10-fold with deionized water) was placed in the test tube. After allowing the mixture to stand at 22 ± 2^°^ C for 5 minutes, 2.5mL of 7.5% Na_2_CO_3_ was added to each combination. The mixture was gently stirred for 20 minutes without allowing any substantial jerking to allow for color development. The UV-Vis spectrophotometer (Model: UV-1700 series) was used to detect the color shift’s intensity at 760 nm. The compound’s TPC was represented in the absorbance value. A final concentration of 0.1 mg/mL was used to appraise the extract and standard samples [18]. Gallic acid was used as the standard to quantify TPCs as mg of GA/g of dry extract, the GA equivalent, or GAE (Standard curve equation: *y* = 0.0085*x* + 0.1125, *R*^2^ = 0.9985), was utilized, represented in Fig 1.

**Fig 1.**
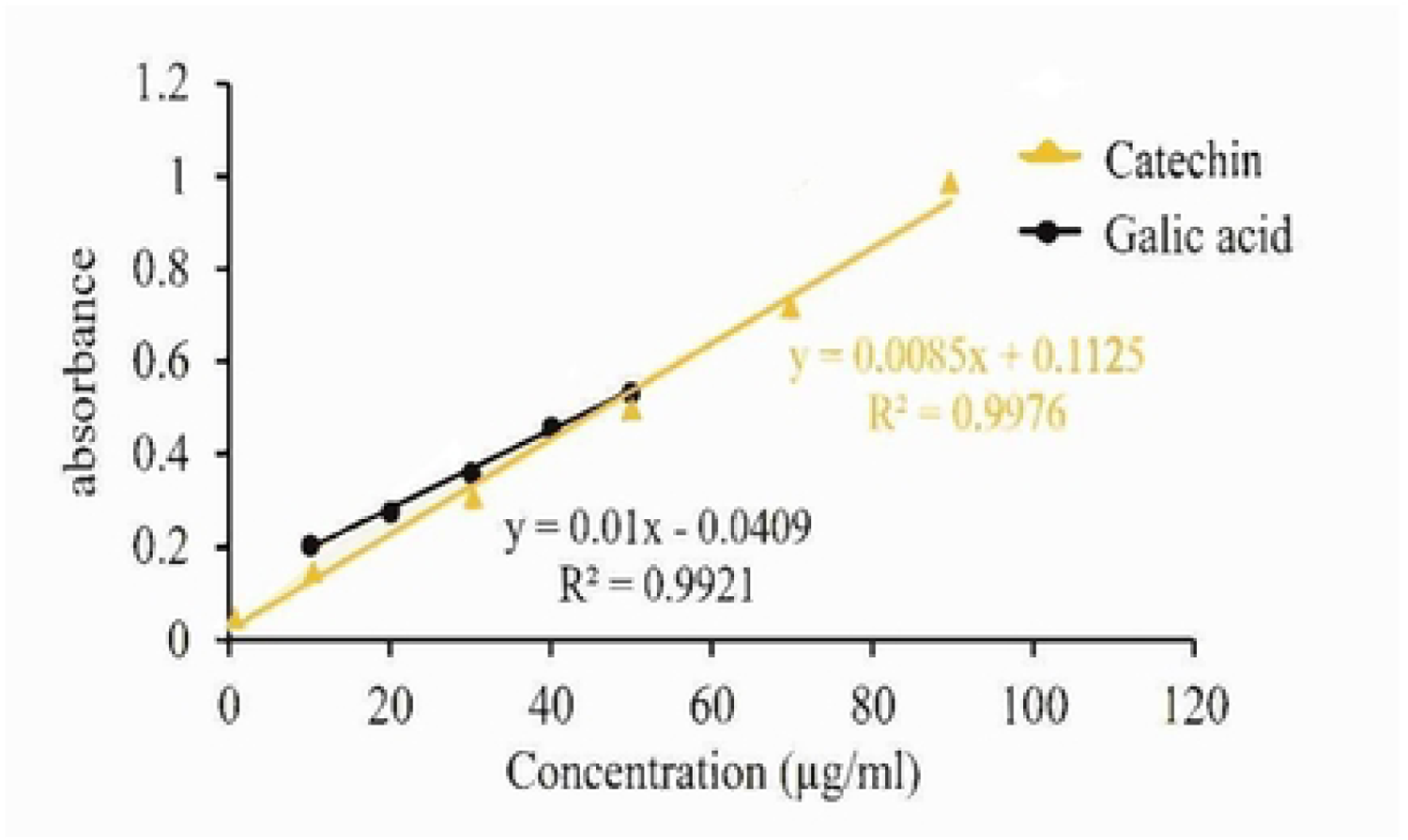
Calibration curve of catechin and gallic acid.

### 2.8 Determination of total flavonoid content (TFC)

The aluminum chloride (AlCl_3_) colorimetric method was utilized to ascertain the total flavonoid content of the crude methanolic extract and its partitionates, which include PSF, MSF, ESF, CTS, and CSF [19]. To summarize, 0.1 mL of 10% AlCl_3_ and 1.5 mL of crude extract (3.12 × 10^-2^ mol/L methanol) were combined, and 0.1 mL of 1 mol/L Na-acetate was then added to the reaction mixture. The mixture was left to stand for half an hour. Next, 1 mL of a 1 mol/L NaOH solution was added, and the combination was finally brought to a final volume of 5 mL using double-distilled water. After the mixture was let to stand for 15 minutes, the absorbance at 415 nm was determined. A calibration curve was used to determine the total flavonoid content. Catechin was used as the standard to quantify TFCs as mg of CE/g of dry extract, the CE equivalent. The total flavonoid content was ascertained using the calibration curve outlined in Fig 1. *Y* = 0.01*X* + 0.0409, *R*^2^ = 0.9921, where X is the catechin equivalent and Y is the crude extract absorbance, represented.

### 2.9 Approval for ethics

International Centre for Diarrheal Disease Research, Bangladesh (ICDDRB) guidelines, which was followed for conducting animal experiments. The World University of Bangladesh Ethical Committee saw the project through and gave its approval (Approval no # WUB/2023/242L6).

### 2.10 Experimental animal

For this experiment, 6-8-week-old Swiss albino mice, weighing 40–45g on average were employed. Mice were kept at a temperature range between 20°C to 24oC. Cages provided with a suitable nesting material to allow mice to regulate their microclimate, in addition to allowing them to perform their natural nesting behaviors. They were given JU-formulated food and water and kept in a typical setting for a week at the World University of Bangladesh research facility in order to help them acclimate

### 2.11 Acute toxicity test

The relatively standard method of Hilaly *et* al. with minor modifications was employed to evaluate the acute toxicity of *Litsea glutinosa* methanol leaves extracts to mice were given different concentrations at oral dosages of 200, 400, 800, 1600, and 3200 mg/kg in order to evaluate the acute toxicity test [20]. Prior to administering extract to each mouse of groups (six mice in each group), were kept for fasting condition till 16 hours. Since all mice were given unrestricted access to food and water, acute toxicity signs were recorded for 48 hours. It should be noted here that a number of fatalities were reported within this time frame [21].

### 2.12 Analgesic activity test

In the current investigation two different methods were employed for testing the possible peripheral and central analgesic effects of *L. glutinosa* leaves; namely acetic acid induced writhing test and hot plate test in mice respectively

### 2.13 The writhing caused by acetic acid

The analgesic activity of the crude methanolic extract of LG leaves was studied using acetic acid induced writhing model in mice. This method previously described by Koster *et* al., [22]. The animals were divided into four groups including negative control (Group I), positive control (Group II) and two test groups (Group III-IV). The animals of group III and IV were administered LG extracts at the dose of 250 and 500 mg/kg body weight respectively. Positive control group received diclofenac (standard drug) at the dose of 25 mg/kg body weight and negative control group was treated with 1% Tween 80 in distilled water at the dose of 10 ml/kg body weight. Test samples, standard drugs and control vehicle were administered orally 30 min before intraperitoneal administration of 0.7% acetic acid. After 15 min of time interval, the writhing (constriction of abdomen, turning of trunk and extension of hind legs) was observed on mice for 5 min. Acetic acid activates the pain nerve and is used to simulate writhing by releasing endogenous chemicals [23]. Each mouse’s writhing number (squirms) was meticulously counted for fifteen minutes.

The percentage inhibition of writhes was calculated using the following formula:

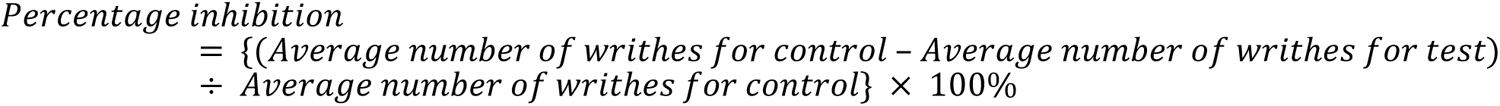

#### 2.13.1 Hot plate test in mice

The method previously described by Eddy and Leimbach was used to screen the extract of LG leaves for centrally acting analgesic activity [24]. The temperature of the hot plate was maintained at 45 ± 0.5⁰C in order to obtain their response to electrical heat-induced nociceptive pain stimulus judged by the presence of behaviors such as licking of the fore and hind paws or jumping. Only animals which responded when placed on the hot plate within a period of 30s were selected for the experiment. Thirty of the selected mice were randomly divided into five groups consisting of six mice each. Groups I and II were orally treated with distilled water (10 ml/kg negative control) and morphine (5 mg/kg positive control) respectively. Groups III and IV were orally pre-treated with crude methanolic extract doses of 250, 500 mg/kg respectively. A cut off time of 20s was used to avoid paw damage. One-hour post treatment, the time taken for each animal to either jump, lick or flutter their paws was taken as the reaction time and recorded at 0, 30, 60, 90 and 120 minutes.

#### 2.13.2 Antipyretic activity test

The antipyretic activity was evaluated by Brewer’s yeast induced pyrexia in experimental animal [25]. Hyperpyrexia was induced by subcutaneous administration of 10 ml/kg body weight 20% aqueous suspension of brewer’s yeast. The selected animals were fasted overnight with water and libitum before the experiments. Initial rectal temperature of animals was recorded using an Ellab thermometer (33.19 ± 0.40°C). After 18h of yeast subcutaneous administration, the animals that showed an increase of 0.3 – 0.5°C in rectal temperature were selected for the antipyretic activity. Crude methanolic extract of plant was given orally at concentration of 100mg, 200mg and 300 mg/kg respectively. Paracetamol (100 mg/kg orally) was used as reference drug, whereas, negative control group received distilled water (10 ml/kg) only. The rectal temperature was recorded up to 3h [26].

#### 2.13.3 Test organisms for antimicrobial studies

The bacteria isolates were obtained from microbiology department of Dhaka University. Bacterial cultures of different strains including *Actinomyces, Bacillus subtilis, Bacillus cereus, Staphylococcus aureus, Sarcina lutea, Salmonella typhi, Chlamydia trachomatis, E. coli, Vibrio mimicus* and *Bacillus parahemolyticus* were maintained at 37°C temperature.

#### 2.13.4 Preparation of inoculum

Five milliliters (mL) of nutritional broth were pipetted into each test tube, corked, autoclaved, and allowed to cool to prepare the bacterial inoculum. The test tubes were labeled and placed on a slide rack. Each test tube was filled with two loopfuls of each test bacteria, thoroughly mixed, and placed in an incubator set to 37°C for 2 hours. Afterward, visible growth of the organisms was observed.

#### 2.13.5 Antibacterial activity

The antibacterial activity of leaves extracts of *L. glutinosa* was assessed using the disc diffusion method. The antimicrobial properties of the carbon tetrachloride fraction (CTF), pet-ether fraction (PSF), ethyl acetate fraction (ESF), methyl soluble fraction (MSF), and chloroform fraction (CSF) were examined and contrasted with a standard antibiotic disc and methanol (solvent was used as negative control in the antibacterial assay). The tests were performed in triplicate and results were recorded as mean ± SD. In summary, agar media was prepared for the culture plates using both gram-positive and gram-negative organisms (*Actinomyces, Bacillus subtilis, Bacillus cereus, Staphylococcus aureus, Sarcina lutea, Salmonella typhi, Chlamydia trachomatis, E. coli, Vibrio mimicus,* and *Bacillus parahemolyticus*).

After being dried and sanitized, a suitable quantity of the test chemicals was impregnated into filter paper discs (6 mm in diameter). Following an evenly seeding of the pathogenic test microorganisms onto each disc, 400 μg of extractives were used to examine the inhibitory action. Nutritious agar medium was used to cultivate five gram-negative and five gram-positive bacterial strains in different dishes. To demonstrate the effectiveness of the antibiotics in conjunction with the test microorganisms and to compare the effects elicited by the known antimicrobial mediator with those elicited by the experimental samples, blank discs were used as the negative control (impregnated with 40 μL methanol solvents). The standard antibiotic Ciprofloxacin (5 g/disc) acted as a positive control. Prior to a 24-hour incubation period at 37°C, the plates were inverted and allowed to diffuse for approximately 24 h at 4°C in a fridge. The diameter of the bacterial growth inhibitory zones was used to quantify the extent to which plant extractives inhibited bacterial growth [27,28].

#### 2.13.6 Statistical evaluation

The experimental data was analyzed using the Statistical Package for the Social Sciences (SPSS) version 22.0 (SPSS Inc., Chicago, IL, USA). The results were determined by taking the average of three evaluations and displaying them as mean ± standard deviation (SD). Significant deviations (p-values < 0.05) between the means were determined by using the using the one-way ANOVA Duncan’s Multiple Range test (DMRT) as post-hoc.

## 3. Results

### 3.1 Analysis of phytoconstituents in the crude methanol extract of LG leaves

Our results of primary phytochemical screening identified several bioactive compounds, including steroids, tannins, saponins, phenols, alkaloids, flavonoids, and other phytocompounds in the crude methanolic extract of LG leaves. In particular, we observed moderate number of phenols, alkaloids, flavonoids, while reducing sugars and steroids were mildly present or absent are displayed in Table 1. In addition to conventional tests of phytocompound screening, in this study we adopted a robust chromatographic technique HPLC to confirm the bioactive compounds present in the LG leaves extract are presented in Fig 2a, 2b, 2c, 2d.

**Table 1.**
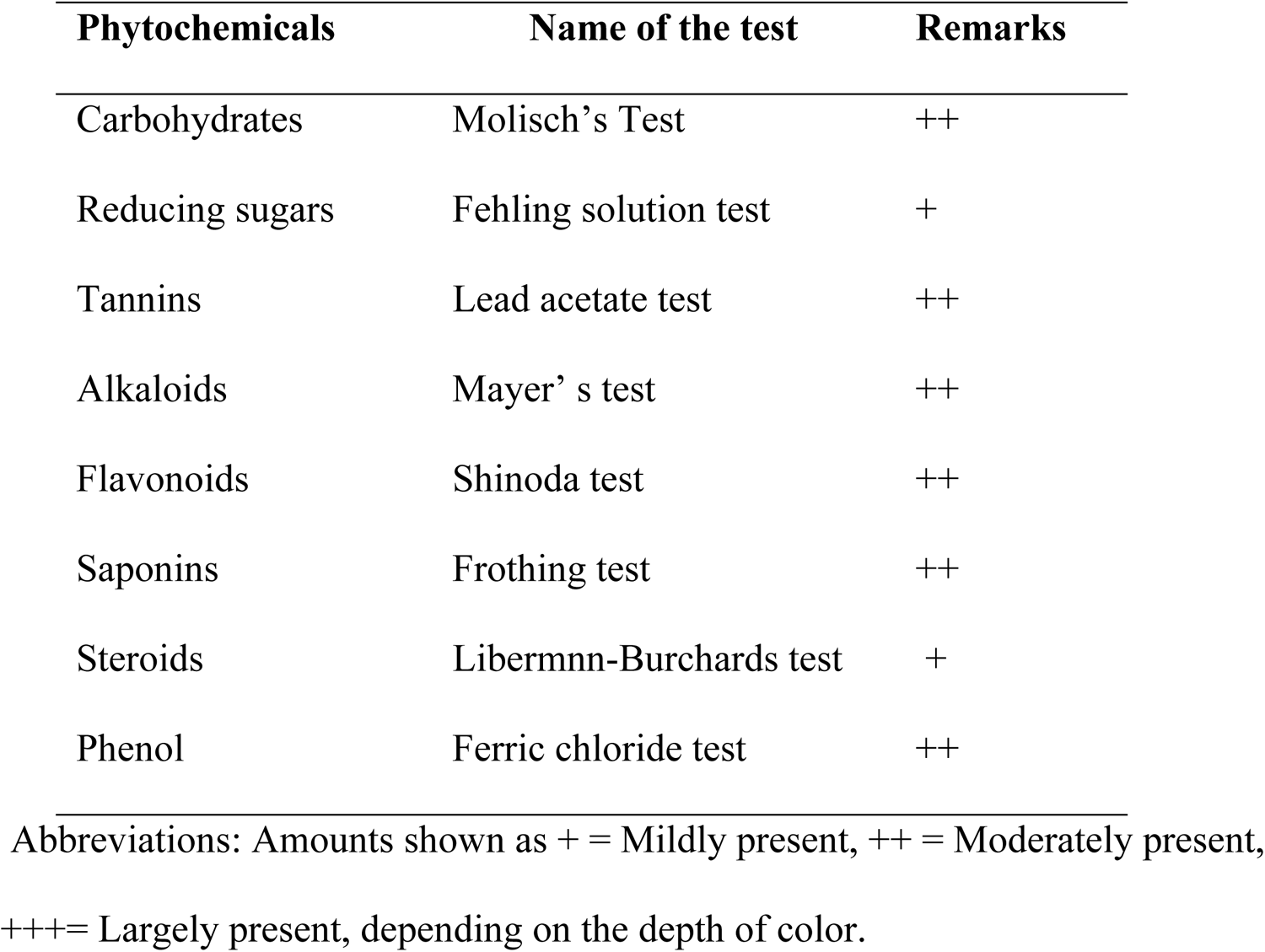
Preliminary phytochemical screening of crude methanolic extract of *Litsea glutinosa* leaves.

**Fig 2a.**
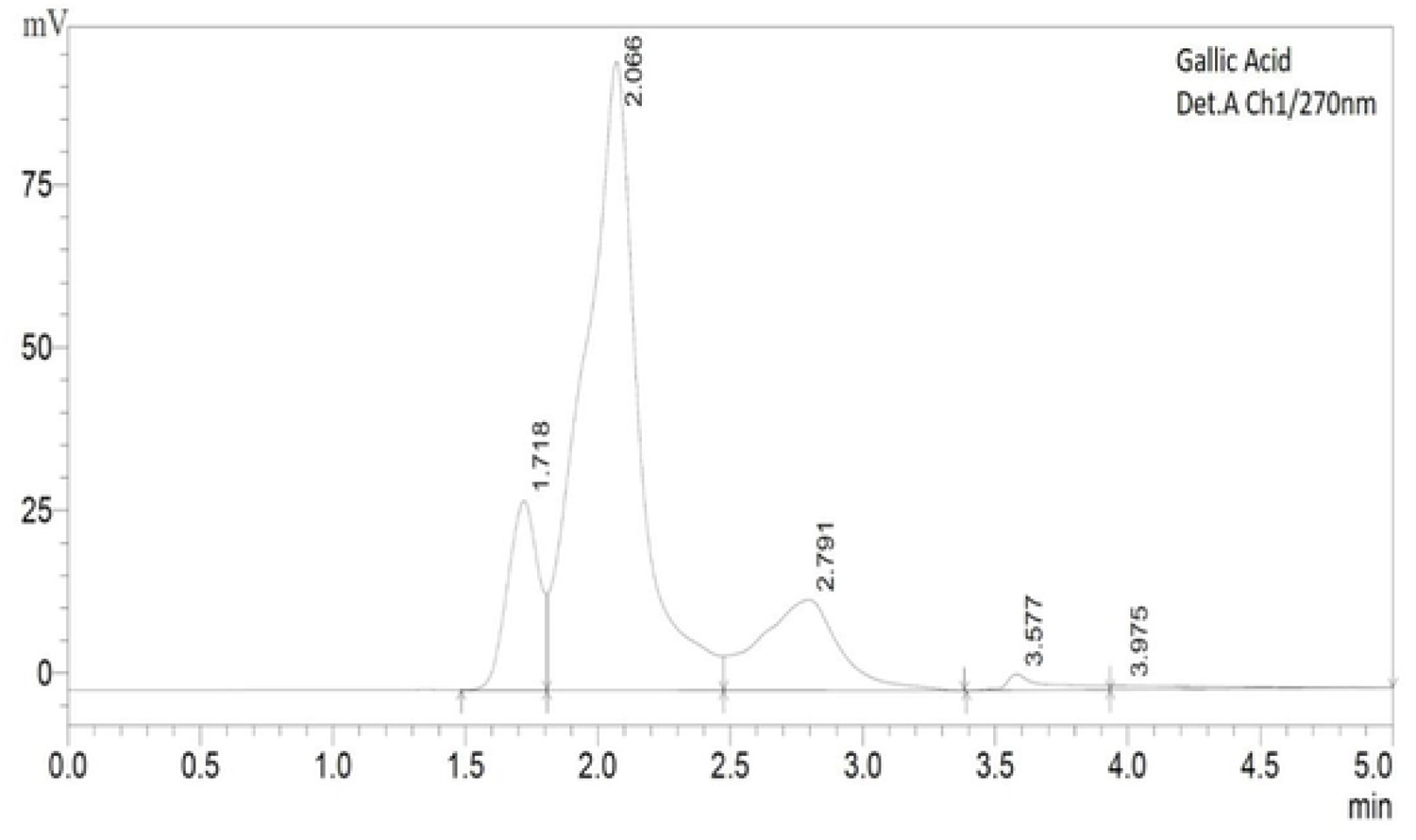
HPLC, Chromatogram for galic acid standard

**Fig 2b.**
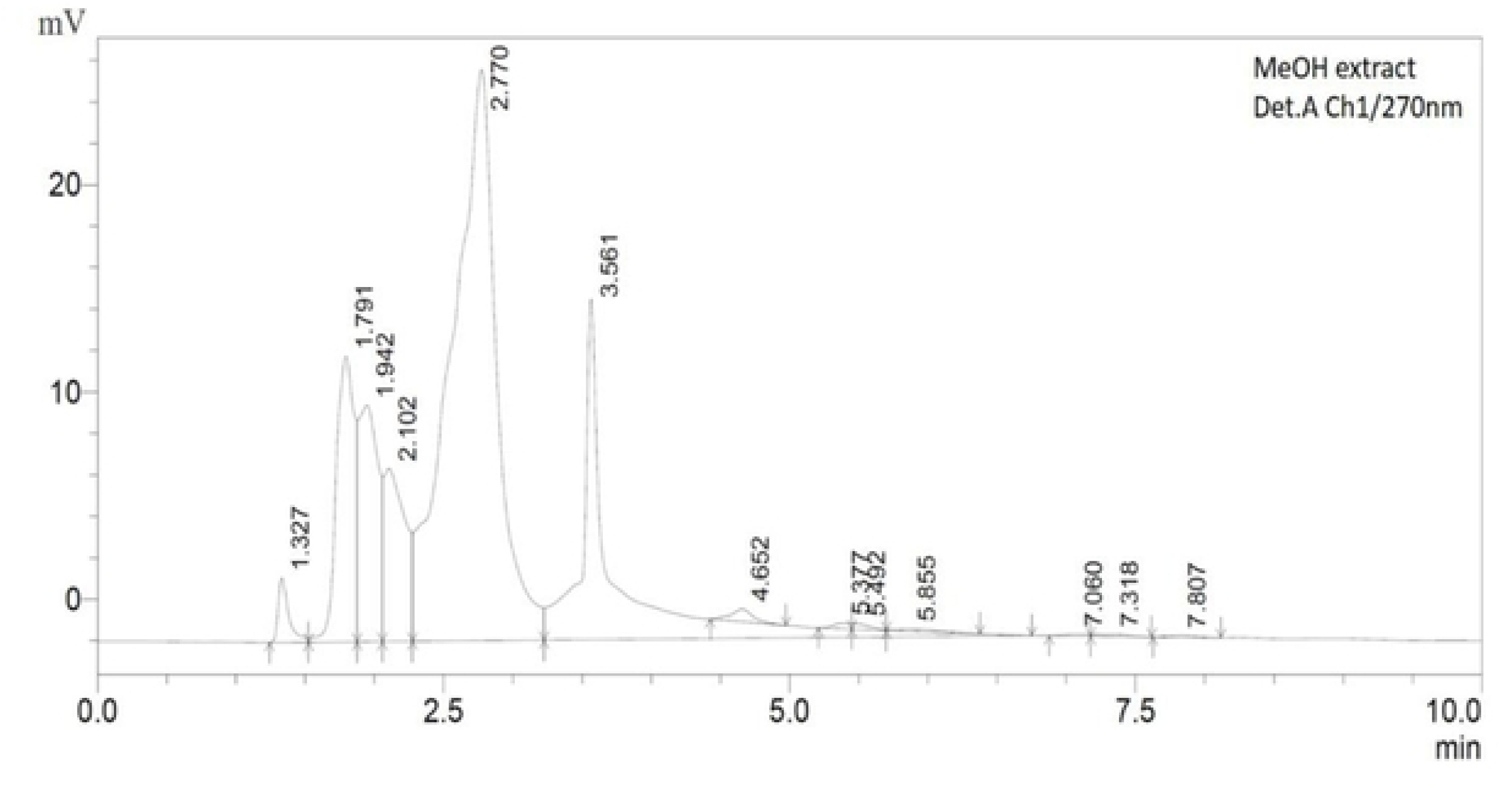
Chromatogram of *Litsea glutinosa* leaves of methanolic extract

**Fig 2c.**
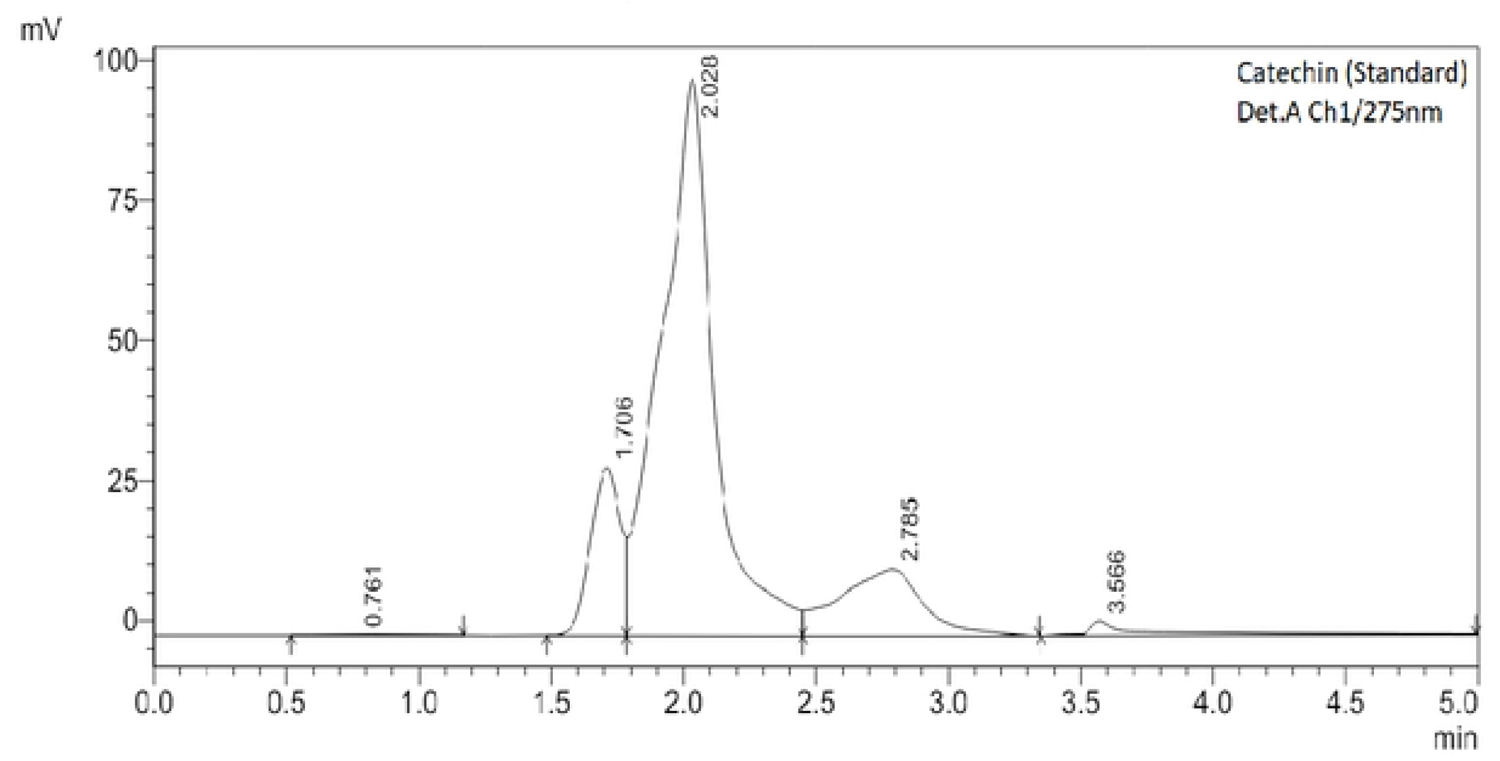
HPLC, Chromatogram for catechin standard

**Fig 2d.**
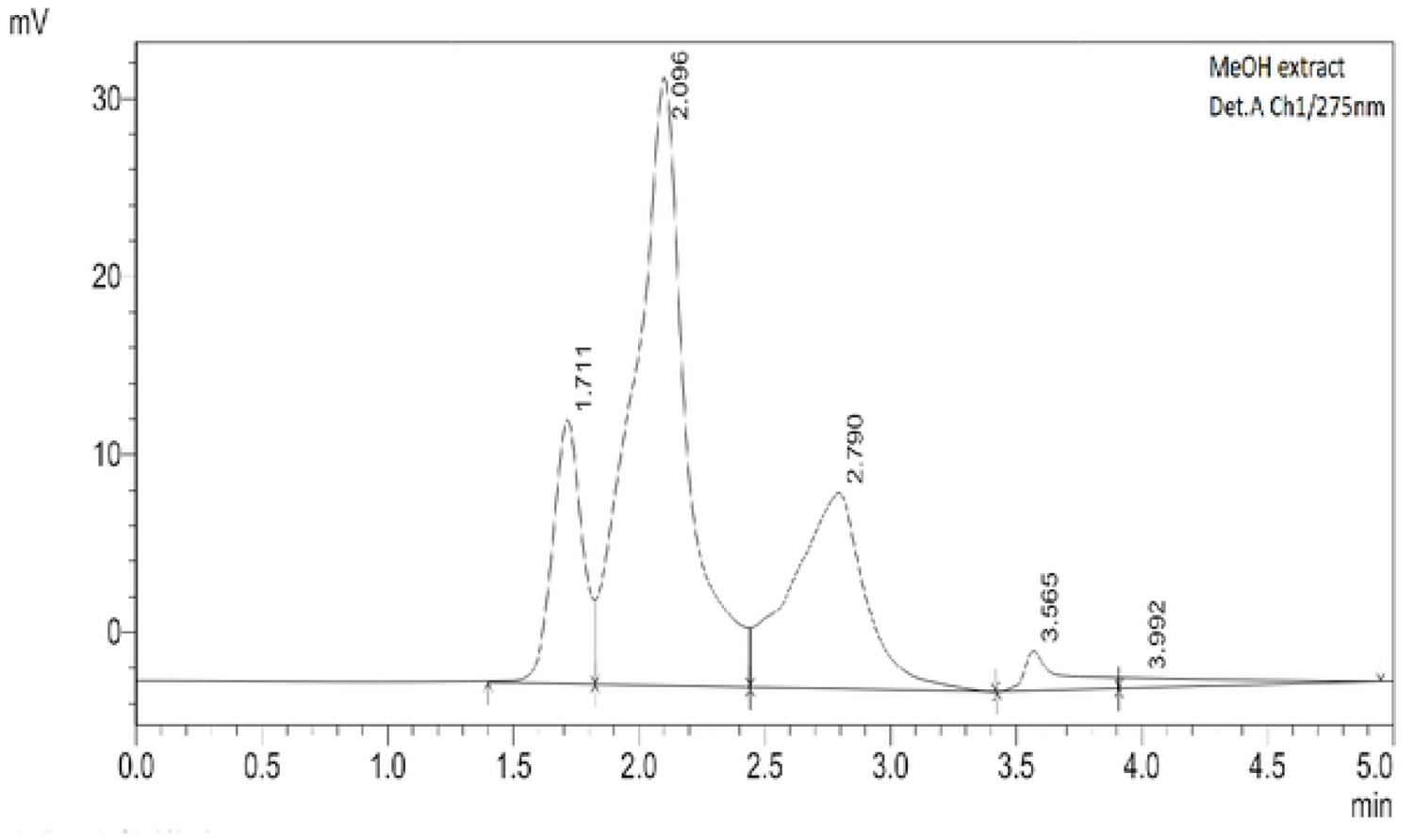
Chromatogram of *Litsea glutinosa leaves* of methanolic extract

### 3.2 Qualitative phytochemical analysis

Several phytochemicals including steroids, tannins, saponins, phenols, alkaloids, flavonoids, and other phytocompounds, were confirmed to exist in the leaves of LG through qualitative investigation shown in Table 1 and quantitative investigation of phenol and flavonoid are presented in Table 2; these phytochemicals may be responsible for producing the analgesic, antipyretic and antibacterial activities.

**Table 2.** Total phenolic and total flavonoid contents of various extractives leave of Litsea glutinosa.

| Sample | TPC (mg of GAE/g of extract) | TFC (mg of CE/g of extract) |
| --- | --- | --- |
| MSF | 135.01 ± 0.57 | 234.30 ± 0.21 |
| PSF | 61.11 ± 0.55 | 131.01 ± 0.33 |
| ESF | 200.61 ± 0.47 | 233.20 ± 0.20 |
| CTF | 110.11 ± 0.37 | 239.31 ± 0.21 |
| CSF | 164.72 ± 0.15 | 290.20 ± 0.12 |

Abbreviations: MSF, Crude methanol extract; PSF, Petroleum ether fraction; ESF, Ethyl acetate fraction; CTS, Carbon tetrachloride fraction; CSF, Chloroform fraction;

In order to confirm the existence of several biologically active phytochemicals, the *Litsea glutinosa* extract and its various organic soluble fractions underwent various tests. Among the investigated extractives were reported dispersible phytochemicals such as gums, tannins, reducing sugars, flavonoids, alkaloids, resins and saponins. It is well recognized in the medical sciences that the phytochemicals acquired in this experiment are beneficial. New therapeutic compounds may be discovered with the help of the current study’s expertise [29].

### 3.3 Chemical standardization of HPLC

HPLC proves to be a highly robust technique for conducting comprehensive analyses of qualitative aspects of phyto compounds within plant extracts. In this study, we used gallic acid as the standard for phenolic compound dentification, while catechin functioned as the standard for identifying flavonoid compounds. The retention time for the phenolic compound standard (gallic acid) was 2.066, and the methanolic extract of LG exhibited a corresponding peak at 2.77 retention time presented in Fig. 2a, 2b. Both the standard and the extract were measured using UVs pectrum at 270 nm, revealing the presence of gallic acid in the methanolic extract, indicating the presence of phenolic compounds. For the flavonoid compound standard (catechin), the retention time was 2.028, with the methanolic extract of *L. glutinosa* displaying a parallel peak at 2.09 retention time displayed in Fig. 2c, 2d. UV spectrum measurements at 275 nm for both the standard and the extract indicated the presence of flavonoids in the methanolic extract. These findings line up with those of previous studies (30,31). This outcome suggests that the methanolic extract of *L. glutinosa* contains both phenolic and flavonoid compounds, which are known for their analgesic, antipyretic and antibacterial activity, and other properties. Consequently, we assessed the total phenolic and flavonoid content in the plant extract was considered a reasonable course of action.

### 3.4 Quantification of total flavonoid content

To assess the catechin equivalent of total flavonoid levels, a complexometric technique employing aluminum chloride was utilized. Using the straight-line *Y* = 0.01 *X* + 0.0409, *R*^2^ ≥ 0.9921 (Fig. 1), which was developed from catechin (with a concentration range of 0 to 160 μg/mL) as the standard, the flavonoid contents were evaluated in terms of catechin equivalent (CE). The flavonoid concentrations of several organic extractives, including MSF, PSF, ESF, CTF and CSF were found to be 234.30 ± 0.21, 131.01 ± 0.33, 233.20 ± 0.20, 239.31 ±0.21, and 290.20 ± 0.12 mg of CE/g of dry extract, respectively presented in Table 2, Fig 3A. The maximum number of flavonoids was observed to be highest in CSF (290.20 ± 0.12 mg of CE/g of dry extract) and lowest in PSF (131.01 ± 0.33mg of CE/g of dry extract), followed by CTF, MSF and ESF. The flavonoid contents of CSF were considerably greater (p < 0.05). The study’s ratification of noteworthy flavonoid concentrations in all partitionates proposes that *Litsea glutinosa* leaves are abundant in flavonoids, which can be used for treating a variety of illnesses. By comparing the results obtained with various solvents, we can show that the extraction values of polyphenols, and flavonoids are greatly depending on the solvent polarity.

**Fig 3A.**
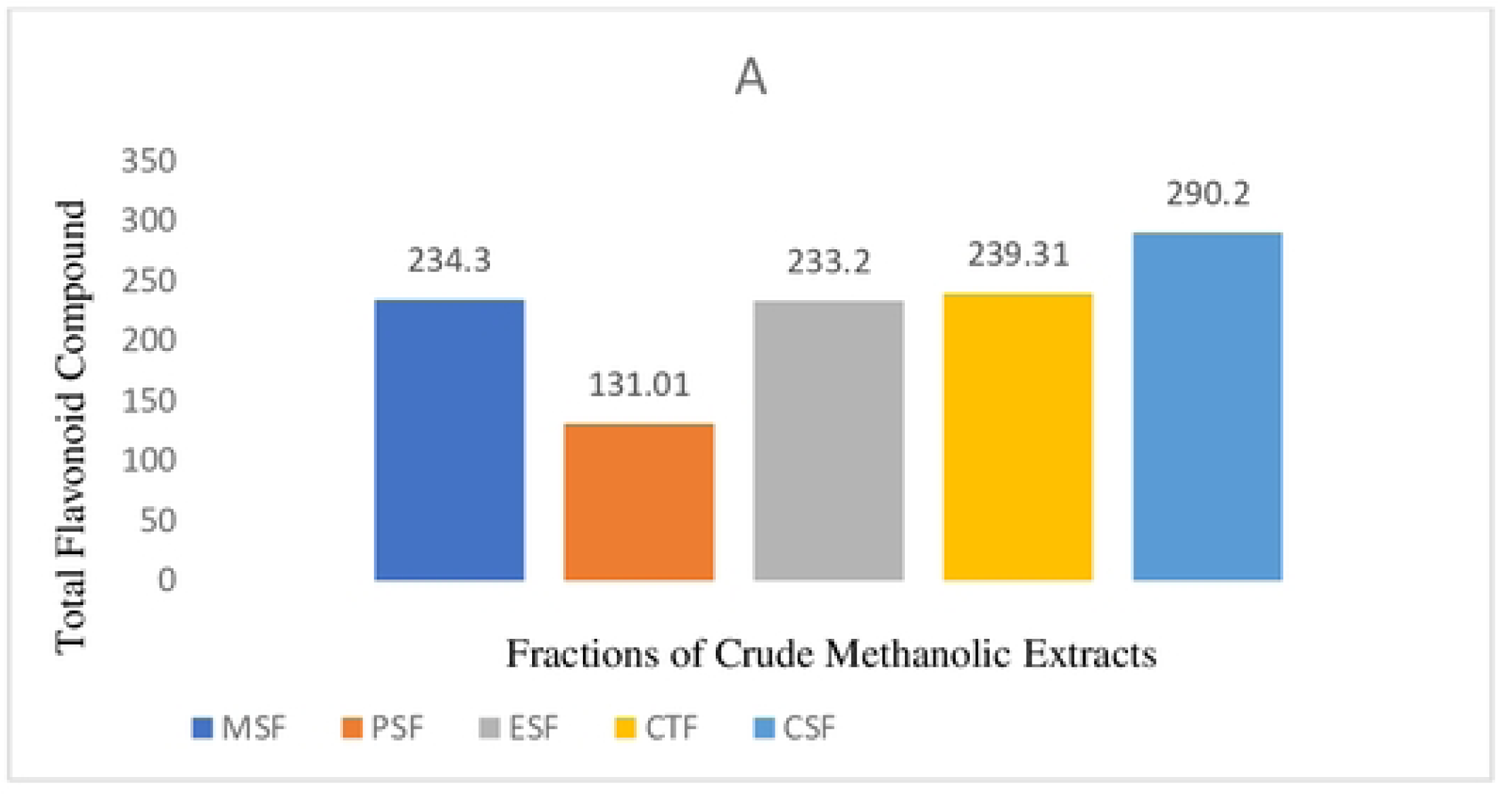
Total flavonoid content study *in vitro* evaluation of fractions of *Litsea glutinosa* leaves

### 3.5 Quantification of total Phenolic contents

The phenolic contents of the crude methanolic extract and its organic soluble fractionates were ascertained using the straight-line *Y* = 0.01 *X* + 0.0501, *R*^2^ ≥ 0.9976 (Fig. 1), with gallic acid (with concentrations ranging up to 20.0 μg/mL) serving as the reference. Phenolic contents of leaves of *Litsea glutinosa extractives* were found to be 135.01 ±0.57, 61.11 ± 0.55, 200.61 ±0.47, 110.11 ± 0.37, and 164.72 ± 0.15 mg of GAE/g of MSF, PSF, ESF, CTF, CSF, in terms of GAE/g displayed in Table 2, Fig 3B. The results indicated that the phenolic contents were highest in ESF (200.61 ± 0.47 mg of GAE/g of dry extract) and lowest in PSF (61.11 ± 0.55 mg of GAE/g of dry extract), with CSF and CTF coming in second and third. In comparison to MSF and PSF, the phenolic contents of ESF were meaningfully higher (p < 0.05). This finding implies that all of the soluble fractions of *Litsea glutinosa* leaves contain phenolic chemicals to varied degrees. The study’s findings indicate that leaves are a significant source of phenolic with possible health advantages.

**Fig 3B.**
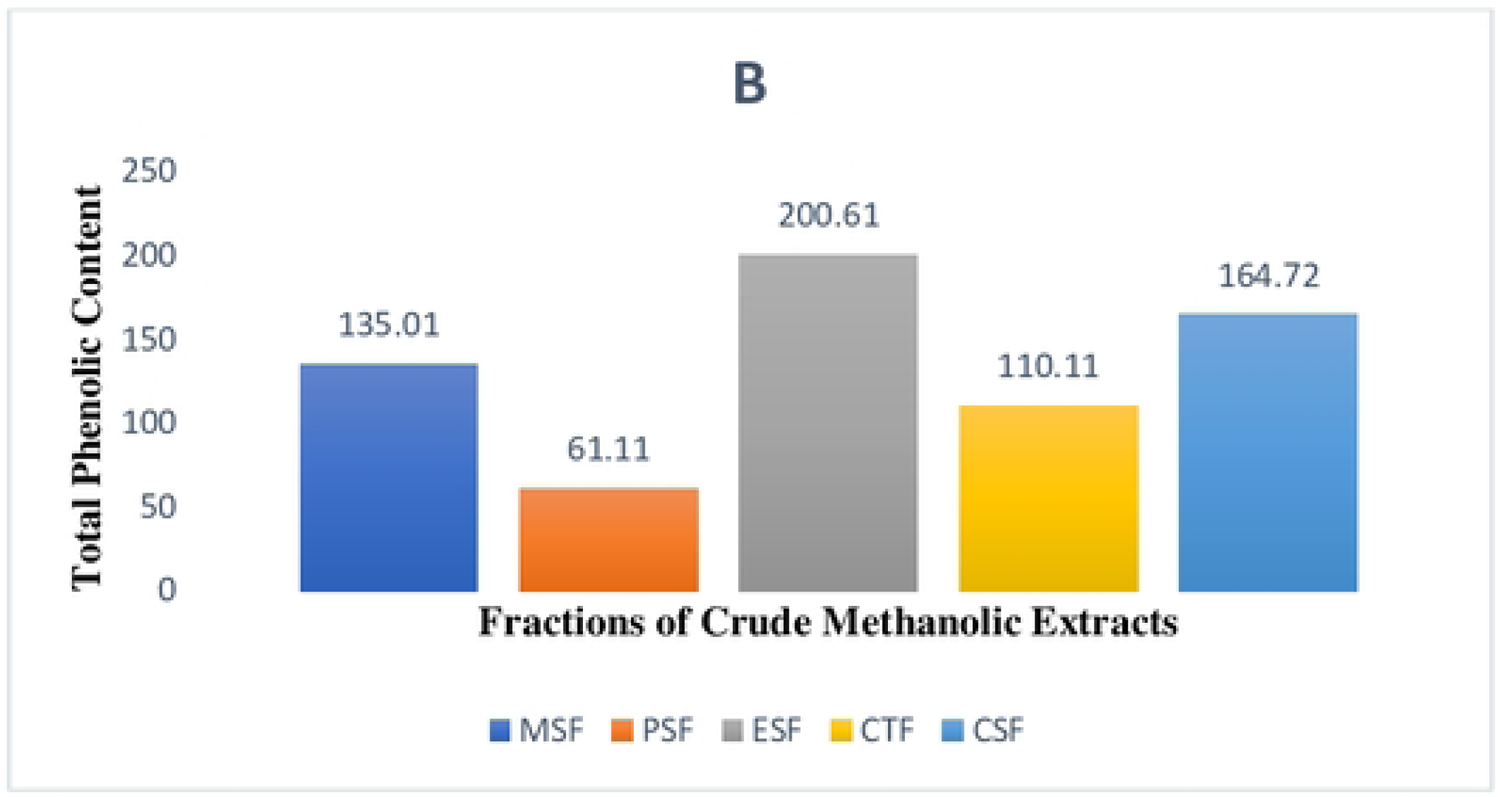
Total phenolic content study of *in vitro* evaluation of fractions of *Litsea glutinosa* leaves

### 3.6 Acute toxicity

No signs of toxicity or deaths were observed after oral administration of oral dosages of 200, 400, 800, 1600, and 3200 mg/kg in order to evaluate the acute toxicity, within the first 48 hours, indicating that the oral LD_50_ is more than above dose. Gross physical and behavioral examinations of the experimental mice revealed no noticeable acute poisoning symptoms, such as vomiting, diarrhea, or loss of appetite.

### 3.7 Analgesic activity (algia)

#### Acetic acid-induced abdominal writhes in mice

While acetic acid was used to prepare the experiment, mice’s writhing was used to test the analgesic activity of the methanolic extract of LG leaves Acetic acid is used to induce writhing and the production of endogenous chemicals that activate the pain nerve. The standard drug diclofenac and the plant extract (test sample) were compared with writing. The writhing response caused by acetic acid is significantly reduced by the LG extract at a dosage of 500 mg/kg body weight. In mice, leaves of LG induced 70% writhing inhibition, compared to 77.5% with standard drug diclofenac at a dosage of 25 mg/kg body weight of the experimental animals as shown in table 3. Values are t-test (n=5); p < 0.001 for standard whereas also p < 0.001 for the leave extract considered as significant % of writhing reduced the number of acetic acid-induced writhing in mice, results showed that the methanolic extract of LG demonstrated a notable analgesic effect. However, more investigation is required to determine the original active component that gives LG’s methanolic extracts their analgesic effects.

**Table 3.** Statistical evaluation of *Litsea glutinosa* leaves extract on acetic acid induced writhing in mice.

| Group | Number of writhing<br>Mean ± SEM | % of Inhibition |
| --- | --- | --- |
| Control (Tween 80) | 32 ± 1.22 | 0 |
| Diclofenac (25 mg/kg) | 7.2 ± 0.84 | 77.5 |
| Plant extract of LG (250 mg/kg) | 10.6±1.00 | 68.1 |
| Plant extract of LG (500 mg/kg) | 9.6 ± 1.14 | 70 |

Abbreviations: Diclofenac as standard drug; LG is as *Litsea glutinosa* leaves and % inhibition = percent inhibition

### 3.8 Hot plate test in mice

At 250 mg ang 500mg methanolic extract of leaves of *L. glutinosa* increased the animal (Swiss-albino mice) reaction time to the thermal stimulus which has been summarized in Table 4. The highest pain inhibition of thermal stimulus was found at a higher dose 500 mg/kg of crude extract which exhibited maximum time for the response against thermal stimuli (3.37± 0.31sec) that is comparable to standard positive control drug morphine (6.47± 0.23a sec) and found statistically significant (P <0.001 and P <0.01) when compared to both the positive control and negative control. Values are presented as Mean ± SEM. Data analyzed using repeated measures ANOVA followed by Bonferroni post hoc test.

**Table 4.**
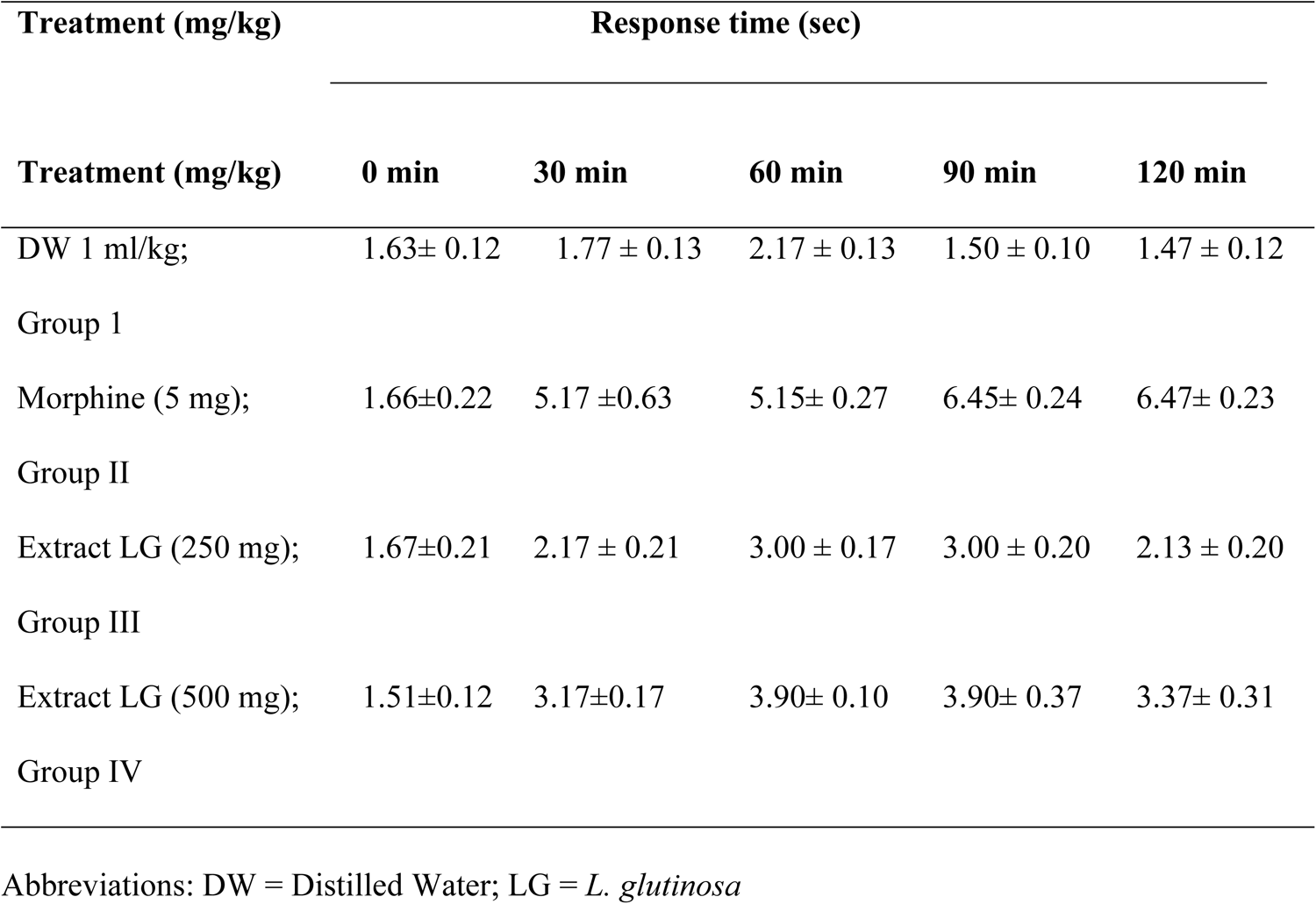
Effect of methanolic leaves extract of *Litsea glutinosa* on hot plate test in mice.

### 3.9 Antipyretic activity of the crude extract

Table 5 showed how several *L. glutinosa* extracts affected the mice’s rectal temperature. Following an 18-hour period, all animals injected with yeast suspension experienced fever, with rectal temperatures varying between 38.03 ± 0.26 and 38.68 ± 0.42 °C. When the standard (at dose 100 mg/kg-body weight) exerted (36.32 ± 0.67°C) after 3 hours of administration, the crude extract demonstrated a significant reduction in body temperature (36.17 ± 0.32°C). The crude extract, at 200 mg/kg, significantly decreased the amount of yeast-induced fever (P < 0.01 at 1, 2, and 3 hours) in comparison to the rectal temperature of the same group at 0 hours. When the 300 mg/kg dose of the crude extract was compared to the same group’s 0-hour rectal temperature, the fever significantly decreased (P < 0.05 at 1, 2, and 3 hours). All dosages of *L. glutinosa* crude extract caused a substantial reduction in rectal temperature (p < 0.05 for doses of 100, 200, and 300 mg/kg, and p < 0.01 for doses of 300 mg/kg) when compared to the control group.

**Table 5.** Effect of the hydroalcoholic extracts of *L. glutinosa* on yeast-induced pyrexia in mice.

| Treatment/Samples | Initial | .5 h | 1 h | 2 h | 3 h |
| --- | --- | --- | --- | --- | --- |
| Control (10 ml/kg) | 38.33 ± 0.44 | 38.70 ± 0.46 | 38.66 ± 0.44 | 38.50 ± 0.56 | 38.39 ± 0.66 |
| Paracetamol standard (100 mg/kg) | 38.28 ± 0.42 | 38.73 ± 0.55 | 38.80 ± 0.68 | 37.50 ± 0.63 | 36.32 ± 0.67 |
| LG Methanolic extract (100 mg/kg) | 38.80 ± 0.30 | 38.17 ± 0.40 | 38.90 ± 0.46 | 37.77 ± 0.25 | 36.60 ± 0.17 |
| LG Methanolic extract (200 mg/kg) | 38.69 ± 0.79 | 38.57 ± 0.25 | 38.27 ± 0.15 | 36.80 ± 0.36 | 36.87 ± 0.32 |
| LG Methanolic extract (300 mg/kg) | 38.43 ± 1.37 | 38.37 ± 0.60 | 38.45 ± 0.70 | 36.00 ± 0.21 | 36.17 ± 0.32 |

Control group received distilled water (10 ml/kg) only; LG as *L. glutinosa*

### 3.10 Antibacterial activity

We tested five organic soluble fractions of LG leaves on five Gram-positive and five Gram-negative bacteria. More pronounce antibacterial action we observed for the extractives against the tested strains *of Actinomyces, Salmonella typhi, E. coli, Bacillus subtilis, Bacillus cereus, Staphylococcus aureus, Sarcina lutea, Shigella dysenteriae, Chlamydia trachomatis* and *Vibrio mimicus,* respectively. Upon observation, it was shown that 400.0 μg of each extractive clearly inhibited the growth of bacteria from 11 to 22 mm displayed in Table 6. The extractives CTF, CSF, and ESF demonstrated notably greater inhibitory zone widths (p < 0.05) against both gram-positive and gram-negative microorganisms.

**Table 6.**
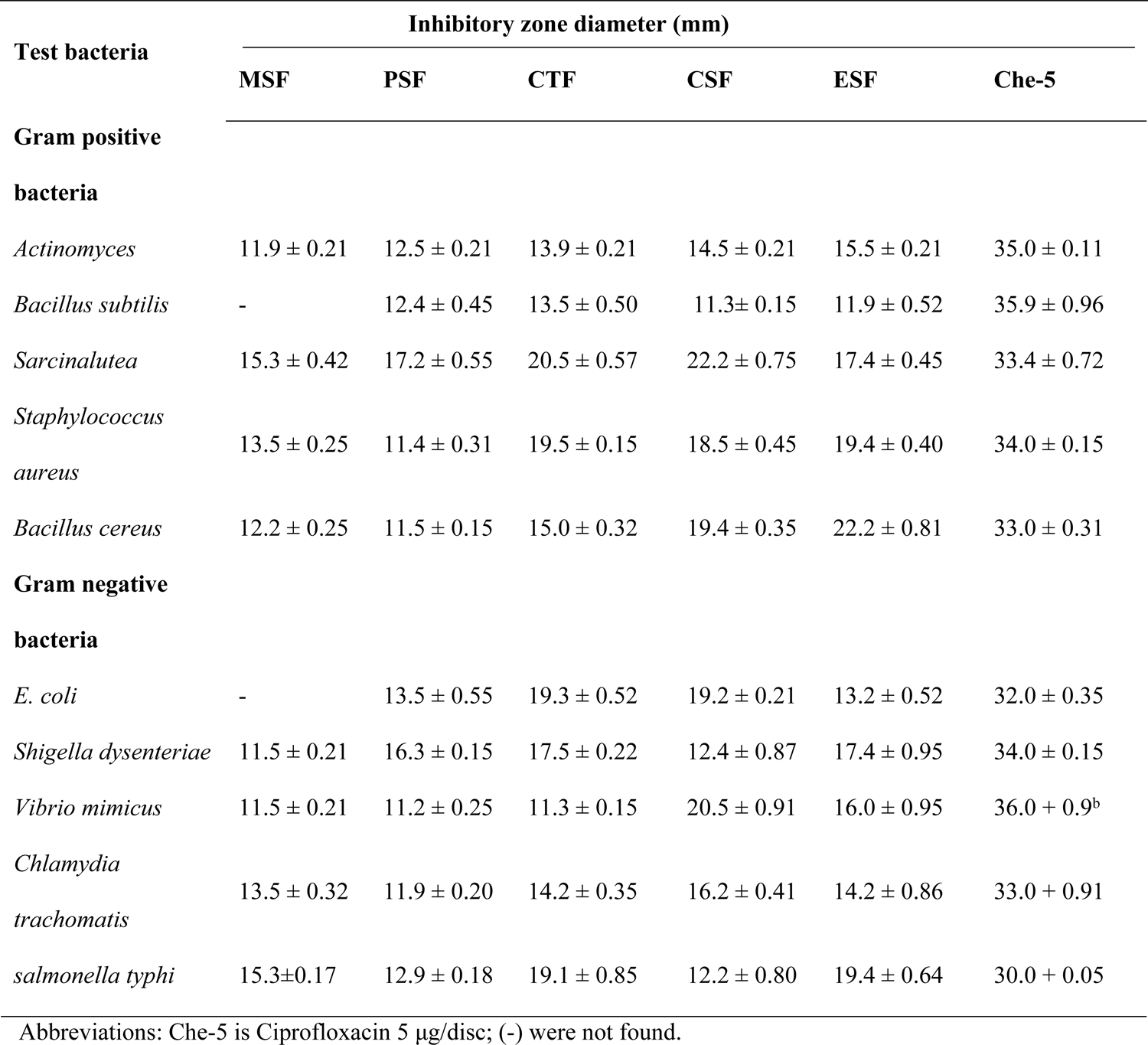
Antibacterial activity by different extractives of leaves of *Litsea glutinosa* through agar diffusion method.

| Test bacteria | Inhibitory zone diameter (mm) |  |  |  |  |  |
| --- | --- | --- | --- | --- | --- | --- |
|  | MSF | PSF | CTF | CSF | ESF | Che-5 |
| **Gram positive bacteria** |  |  |  |  |  |  |
| *Actinomyces* | 11.9 ± 0.21 | 12.5 ± 0.21 | 13.9 ± 0.21 | 14.5 ± 0.21 | 15.5 ± 0.21 | 35.0 ± 0.11 |
| *Bacillus subtilis* | - | 12.4 ± 0.45 | 13.5 ± 0.50 | 11.3 ± 0.15 | 11.9 ± 0.52 | 35.9 ± 0.96 |
| *Sarcinalutea* | 15.3 ± 0.42 | 17.2 ± 0.55 | 20.5 ± 0.57 | 22.2 ± 0.75 | 17.4 ± 0.45 | 33.4 ± 0.72 |
| *Staphylococcus aureus* | 13.5 ± 0.25 | 11.4 ± 0.31 | 19.5 ± 0.15 | 18.5 ± 0.45 | 19.4 ± 0.40 | 34.0 ± 0.15 |
| *Bacillus cereus* | 12.2 ± 0.25 | 11.5 ± 0.15 | 15.0 ± 0.32 | 19.4 ± 0.35 | 22.2 ± 0.81 | 33.0 ± 0.31 |
| **Gram negative bacteria** |  |  |  |  |  |  |
| *E. coli* | - | 13.5 ± 0.55 | 19.3 ± 0.52 | 19.2 ± 0.21 | 13.2 ± 0.52 | 32.0 ± 0.35 |
| *Shigella dysenteriae* | 11.5 ± 0.21 | 16.3 ± 0.15 | 17.5 ± 0.22 | 12.4 ± 0.87 | 17.4 ± 0.95 | 34.0 ± 0.15 |
| *Vibrio mimicus* | 11.5 ± 0.21 | 11.2 ± 0.25 | 11.3 ± 0.15 | 20.5 ± 0.91 | 16.0 ± 0.95 | 36.0 ± 0.9<sup>b</sup> |
| *Chlamydia trachomatis* | 13.5 ± 0.32 | 11.9 ± 0.20 | 14.2 ± 0.35 | 16.2 ± 0.41 | 14.2 ± 0.86 | 33.0 ± 0.91 |
| *salmonella typhi* | 15.3 ± 0.17 | 12.9 ± 0.18 | 19.1 ± 0.85 | 12.2 ± 0.80 | 19.4 ± 0.64 | 30.0 ± 0.05 |

Abbreviations: Che-5 is Ciprofloxacin 5 µg/disc; (-) were not found.

## 4 Discussion

Polyphenols, including flavonoids, phenolic acids, tannins, lignans, and coumarins, are naturally occurring phytochemicals found in fruits, vegetables, cereals, roots, and leaves. Recent research highlights the potential health benefits of these compounds, particularly as antioxidants combating oxidative stress-related diseases. Phenolic compounds have garnered significant attention due to their antioxidant, antimicrobial, anti-inflammatory, anticancer, and cardiovascular protective activities [32]. Our study’s quantitative analysis confirms that naturally occurring phytochemical compounds in *Litsea glutinosa* leaves contain satisfactory quantities of phenols and flavonoids, encouraging further evaluation of their pharmacological potential.

In the acetic acid-induced writhing test for peripheral analgesia in mice, pretreatment with 500 mg/kg b.w. of the ethanolic extract significantly reduced the writhing frequency by 70% (p < 0.001), compared to 77.5% for the standard diclofenac 25mg. Besides peripheral nociception, the LG extract also suppressed the central nervous pain response by significantly increasing the latencies and basal pain thresholds in the hot plate tests. The acetic acid-induced abdominal writhing test effectively screens compounds for peripherally and centrally acting analgesic activities [35]. Following intraperitoneal injection of acetic acid, endogenous chemicals such as PGE2, PGF2α, PGI2, serotonin, histamine, lipoxygenase products, and peritoneal mast cells are released and accumulated, directly activating nociceptors and causing pain [33]. Mice react to this chemical stimulus by contracting their abdominal muscles, elongating their body parts, and extending their rear limbs. This response is believed to be mediated by local peritoneal receptors [34].

Aspirin and other NSAIDs reduce acetic acid-induced writhing by delaying the synthesis or release of these endogenous pain and inflammatory mediators [35]. The reduction in acetic acid-induced writhing by the leaves extract suggests peripherally mediated analgesic activity through blocking the synthesis or release of endogenous substances responsible for pain sensations. In the hot plate method, the extract significantly delayed the mice’s reaction latency to thermally generated pain. However, morphine, the conventional medication, more effectively inhibited the animals’ thermal pain response than the leaves extract. Centrally acting analgesic compounds delay the response time to a heat stimulus [36]. Therefore, the leaves extract’s ability to extend the latency period for pain suggests activity via central pain pathways, likely by modulating endogenous substances that target pain and inflammation, in addition to opioid receptors, such as endogenous opioids, somatostatin, and other inhibitory hormones [37].

Fever in animal models is often induced by exogenous substances, including bacterial endotoxins and microbial infections. These exogenous pyrogens trigger the production of various pro-inflammatory cytokines, which stimulate the release of local prostaglandins (PGs) that enter the hypothalamic circulation and reset the hypothalamic thermal set point. Maintaining body temperature requires a balance between heat production and loss, regulated by the hypothalamus. Non-steroidal anti-inflammatory drugs (NSAIDs) demonstrate antipyretic action by inhibiting prostaglandin synthetase within the hypothalamus [38]. Our study revealed that the methanolic extract of L. glutinosa leaves significantly reduces yeast-induced elevation of body temperature, producing effects similar to the standard drug. It is hypothesized that the extract exerts its antipyretic action by inhibiting prostaglandin synthetase within the hypothalamus, akin to NSAIDs. Although direct evidence of *L. glutinosa* interfering with hypothalamic prostaglandin synthesis is lacking, a related study found that Dalbergia odorifera extract inhibits prostaglandin biosynthesis [39]. Thus, the present pharmacological evidence supports the folklore claim of L. glutinosa leaves as an antipyretic agent.

In antibacterial studies, results demonstrated that a 400.0 μg dose of each extract inhibited bacterial growth, with inhibition zones ranging from 11 to 22 mm. The extractives CTF, CSF, and ESF showed significantly larger inhibitory zones (p < 0.05) against tested strains compared to the standard antibiotic ciprofloxacin 5.0 μg/disc. Ciprofloxacin, a second-generation fluoroquinolone, is effective against many Gram-negative and Gram-positive bacteria by inhibiting bacterial DNA gyrase and topoisomerase, binding to bacterial DNA gyrase with 100 times the affinity of mammalian DNA gyrase [40]. The current results suggest that the LG extract and its fractions exhibit antibacterial activity, likely through inhibition of bacterial DNA gyrase and topoisomerase. Previous reports indicate that various organic solvent extracts of *L. glutinosa* leaves are effective against a range of Gram-positive and Gram-negative bacteria. Plant polyphenols and flavonoids with antioxidant activity have been shown to exhibit potential antibacterial activity [41,42]. This study examined the antibacterial efficacies of extracts against nine different bacteria, confirming that *L. glutinosa* leaves extract exhibits substantial antibacterial activity

## 5 Conclusion

The present study employed phytochemical screening to identify bioactive compounds in the methanolic crude extracts of *L. glutinosa* leaves and their various soluble fractionates, for instance, flavonoids, reducing sugars, tannins, gums, saponins, quinines, glycosides, steroids, and terpenoids. Moreover, the plant extracts demonstrated significant analgesic, antipyretic and antibacterial activities, Thus, the current research justifies the application of *L. glutinosa* in traditional medicine to treat a range of ailments. Isolating the active chemical components in charge of the pharmacological qualities will require more investigation. These findings provide light on the enormous potential of natural resources to promote human health, which may pave the way for the development of innovative drugs and nutraceuticals in the future. Nevertheless, more investigation, encompassing clinical trials, is imperative to corroborate and convert these encouraging outcomes into concrete medicinal uses.

## Acknowledgement

The author gratefully acknowledges with sincere thanks, Department of Pharmacy of the World University of Bangladesh provided complete laboratory facilities for this research project. Each author has approved submission and taken full responsibility for the content of the work that has been submitted.

## Author Contributions

Conceptualization: Zubair Khalid Labu, Md. Imran Hossain Data curation: Md. Shakil Formal analysis: Zubair Khalid Labu, Samira Karim Investigation: Samira Karim, Md. Tarekur Rahman Methodology: Tarekur Rahman, Zubair Khalid Labu Project administration: Samira Karim, Md. Tarekur Rahman Supervision: Zubair Khalid Labu, Md. Imran Hossain Writing – original draft: Zubair Khalid Labu, Md. Imran Hossain Writing – review & editing: Tarekur Rahman, Zubair Khalid Labu

